# Culture of preimplantation embryos in media containing L-proline increases intracellular GSH concentration throughout development

**DOI:** 10.64898/2026.04.23.720483

**Authors:** Madeleine L.M. Hardy, Michael B. Morris, Margot. L Day

**Affiliations:** School of Medical Sciences, Faculty of Medicine and Health, The University of Sydney, NSW 2006, Australia

## Abstract

Careful balance of the redox status of the embryo and reduction of oxidative stress is crucial in early development. Here we show that the culture of preimplantation mouse embryos in the conditionally non-essential amino acid L-proline (Pro) increases the intracellular concentration of the potent antioxidant glutathione as shown by staining of 2-cell, 4-cell and 8-cell embryos with tetrafluoroterephthalonitrile (4F-2CN). Further, liquid-chromatography/mass spectrometry showed increased GSH levels in all Pro-treated preimplantation stages of development compared to controls,. The GSH:GSSG ratio also showed a Pro-dependent increase. Overall, our results indicate that the beneficial effect of Pro in preimplantation embryo culture is due to the reduction in oxidative stress mediated through an increase in cellular GSH concentration.

## Introduction

The tripeptide glutathione (GSH; γ-L-glutamyl-L-cysteinylglycine) is the major thiol antioxidant in mammalian cells and regulates several key cellular processes through its antioxidant function [1]. Cellular concentrations of GSH range from 1-20 mM, with concentrations of the oxidised form, GSSG, typically substantially lower (in the micro to nanomolar ranges) [2]. GSH protects cells from oxidative distress, which if left uncorrected lead to a variety of cellular dysfunctions, including cell death and failure to progress through the cell cycle [3,4]. GSH also modulates cell signalling [5] by reducing selected, often proteinaceous, Cys dilsulphide bridges and reversibly binding to thiol groups in a process known as glutathionylation [6].

Glutathione synthesis and metabolism occur through a series of enzymatic reactions in the mitochondria and cytoplasm [7]. Figure 1 shows: (i) The stepwise formation of GSH from its component amino acids (AAs) centring on Pro-mediated synthesis. (ii) The GSH redox cycle, whose major role is to reduce cellular concentration of highly reactive (and potentially damaging) H_2_O_2_ to homeostatic concentrations (1–10 nM) and to generate NADP^+^, used principally for the production of NADPH for the regulation of cellular metabolism and maintaining redox homeostasis [8].

**Figure 1.**
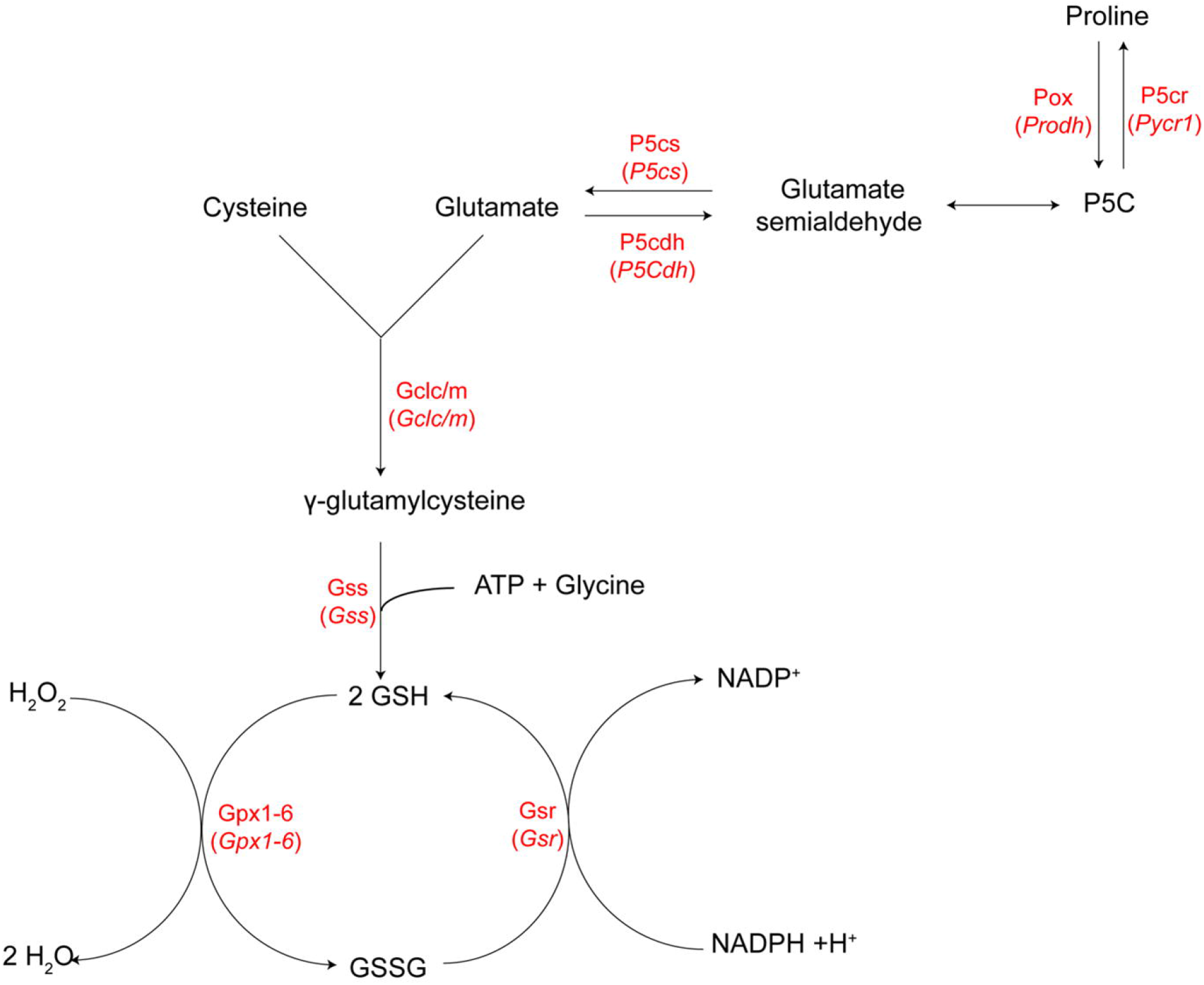
GSH synthesis and redox cycle. Pro can be metabolised to 1-pyrroline-5-carboxylate (P5C) in the proline cycle by proline oxidase (*Prodh*). P5C then spontaneously metabolises to glutamate semialdehyde which can be enzymatically converted to Glutamate. The tripeptide GSH is formed initially by ligation of cysteine and glutamate by γ-glutamylcysteine ligase (*Gcl*) which has both a catalytic (*Gclc*) and modifying (*Gclm*) subunit. This is followed by the ATP-dependent addition of glycine by glutathione synthetase (*Gss*). The resulting GSH enters a redox cycle in which GSH is converted to its oxidised, dimeric form, GSSG, by glutathione peroxidase (*Gpx*) enzymes 1-6, while simultaneously reducing highly reactive H_2_O_2_ to H_2_O. GSSG is reduced back to GSH by glutathione reductase (*Gsr*) with the help of the cofactor NADPH, thereby producing NADP^+^ for use in redox homeostasis and cellular metabolism. Enzymes are in red. Genes encoding these enzymes are italicised in parentheses.

Oxidative stress is controlled in preimplantation embryos and oocytes through intracellular redox mechanisms and antioxidant systems present in the follicular, oviductal and uterine fluids [9]. This extracellular regulatory control is largely absent in oocyte/embryo culture conditions resulting in increased levels of reactive oxygen species (ROS) and poorer development [9,10].

Intracellular GSH concentrations as well as the extracellular concentrations in reproductive fluids correlate with developmental success [9]. Follicular fluid contains antioxidants such as GSH and Cys, as well as enzymes related to their synthesis, including Gpx, Gsr and Gss [11]. Follicular fluid collected from women undergoing IVF treatment contains 8-10 µM GSH [12,13], while mouse oviductal fluid contains approximately 60 nmol GSH/mg protein, and uterine secretions between 60-120 nmol GSH/mg protein, depending on the stage of the oestrous cycle [9,14].

Intracellular GSH concentration in the oocyte is critical during fertilisation *in vivo* where it is required to compensate for the high levels of ROS the oocyte is exposed to by the sperm, and to allow for sperm nuclear condensation [15]. As the zygote develops into the blastocyst *in vivo*, GSH levels decrease to less than 10% of that seen in the oocyte [16].

During the earliest stages of embryo cleavage intracellular GSH levels are very low but the presence of other antioxidants and antioxidant enzymes, including superoxide dismutase and catalase, compensate for this to maintain redox balance [14,17,18]. However, GSH is still important in these early cleavage stages since when GSH concentrations are reduced by 90% in 2-cell embryos using diethyl maleate (DEM), there is a dose-depended decrease in morula and blastocyst formation [14]. Blastocysts depleted of GSH using buthionine sulfoximine (BSO) recover their GSH stores when BSO is removed, faster than 2-cell embryos. This is due to the fact the production of GSH is increased throughout preimplantation development due to the increase in activity of GSH synthesis enzymes [14].

In the early embryo, GSH production reduces hydrogen peroxide [19](Figure 1), which in turn prevents histone methylation errors and DNA damage [20].

Cysteine is the rate limiting component of GSH production [21]. In the oocyte and early embryo, prior to zygotic genome activation, SLC1A5 (Asct2) or SLC7A11 (xCT^−^) transporters promote extracellular cysteine uptake [22,23]. Beyond this stage, cysteine is produced by metabolism of other amino acids including methionine and serine [22]. Exogenous addition of cysteine to IVM media improves maturation of caprine and bovine oocytes, although this result is contentious with some studies finding there is no effect on maturation [24–26]. This is likely to due to a combination of factors including increased production of GSH and subsequent reduction of ROS as well as increase in NADPH availability and homocysteine production [22]. Have to find some way to round out this para,

The non-essential AA proline (Pro) improves preimplantation embryo development when added to culture medium [27]. When added to IVF medium, Pro reduces intracellular ROS concentration and mitochondrial activity by 60% and 40%, respectively, and subsequent development to the blastocyst stage in the absence of Pro increases from 35% to 60%, with the effect observed from compaction onwards [28]. Pro similarly reduces ROS and mitochondrial activity in 2-cell and 4-cell embryos when included in culture medium from the zygote stage [29].

In various systems, prolonged exposure to Pro can reduce ROS levels and/or mitochondrial activity by a variety of mechanisms [17,28,29]. One mechanism for reducing ROS levels is by increasing GSH concentration via Pro’s metabolism to Glu (Figure 1) but this has not been shown in preimplantation embryo culture. However, Pro-mediated increases in GSH concentration does occur in several cell types, including mouse oocytes [30], boar sperm [31], HEK293 cells and porcine trophectoderm cells [32,33] suggesting this as a mechanism for improving preimplantation development.

Tetrahydro-2-furoic acid (THFA) is an inhibitor of proline oxidase (POX) which metabolises Pro to pyrroline-5-carboxylate (P5C) which is the first step in the metabolism of Pro [34,35]. When THFA is included in embryo culture media it prevents the Pro mediated improvement in development reducing the percentage of embryos that develop to the blastocyst. Further, when embryos are cultured to the 2-cell or 4-cell stage with 400 µM THFA + 400 µM Pro, Pro is unable to reduce ROS levels or mitochondrial activity [29].

The hypothesis for this study is that one of the mechanisms by which Pro improves preimplantation embryo development is by increasing the concentration of GSH and reducing oxidative stress. To test this, the effect of embryo culture in the presence of Pro on GSH and Cys levels was examined in 2-cell, 4-cell and 8-cell mouse embryos using fluorescence probes and confocal microscopy. The effect of Pro on (i) GSH and GSSG concentrations (measured using LC/MS) and (ii) expression of genes involved in conversion of Pro to GSH (by qPCR) was examined.

## Materials and methods

### Animals (*Mus musculus)*

Experiments were performed using outbred Quackenbush Swiss (QS) mice (Animal Research Centre, Perth Australia and Lab Animal Services, The University of Sydney). Animals were used in accordance with the Australian Code of Practice for the Care and Use of Animals for Scientific Purposes and was approved by the University of Sydney Animal Ethics Committee as per the requirements of the NSW Research Act (approval numbers 842 and 1877).

### Zygote collection and media preparation

Superovulation was induced in 3-5 week-old mice using intraperitoneal injection with 10 IU pregnant mares’ serum gonadotrophin (PMS) (Intervet, Vic, Australia) followed 48 h later by injection of 10 IU human chorionic gonadotropin (hCG) (Intervet, Vic, Australia). Females were paired with 2-8 month-old males overnight for mating. Females were euthanised by cervical dislocation 24 h post hCG and the oviducts isolated. Zygotes were dissected from the oviducts in HEPES buffered modified tubal fluid (HEPES-ModHTF) [27] and treated with hyaluronidase (1 mg/mL in HEPES-ModHTF) to remove the cumulus cells. Zygotes were then transferred to modified human tubal fluid (ModHTF) without Gln. Media were supplemented with 0.3 mg/mL bovine serum albumin (BSA), buffered to pH 7.4 and adjusted to 270 mOsm/kg.

### Live-cell fluorescence imaging of glutathione and cysteine

Zygotes were cultured at high density (1 embryo / 1 µL) in ModHTF (modified human-tubal fluid) overlayed with mineral oil. Zygotes were cultured to the 2-cell, 4-cell, or 8-cell stage for 48, 60 or 76 h respectively, in ModHTF medium containing no AAs or containing 400 µM Pro, tetrahydro-2-furoic acid (THFA) or Pro + THFA at 37 °C and 5% CO_2_.

Prior to staining, 3-5 embryos that had been cultured in the absence of Pro and or THFA were exposed to 0.25 µM Cys to increase Cys concentrations as a positive control or exposed to 2 mM N-ethylaldemide (NEM) to reduce GSH fluorescence. NEM acts as a thiol depletion agent sequestering active GSH to GS-NEM and subsequently preventing GSH fluorescence.

Tetrafluoroterephthalonitrile (4F-2CN) is a simple molecule that acts as a fluorescence probe by undergoing selective reactions with between the cyano group in the probe and the thiols in GSH and cysteine. The thiol groups are transformed by the probe and can be differentially excited at different wavelengths [36]. 4F-2CN emits strong blue fluorescence when it reacts with GSH and strong green fluorescence when it reacts with Cys.

Embryos were transferred in groups of up to 10 to wells containing ModHTF in a 96-well plate with each treatment and 20 µM 4F-2CN (Sigma Aldrich, #104426) to stain for GSH and Cys. Embryos were incubated in medium containing the dye for 30 min at 37 °C and 5% CO_2_. Embryos were then washed twice in ModHTF containing Pro, THFA or Pro +THFA to remove excess stain. A 35 mm glass-bottomed petri dish was prepared containing 10 μL drops ModHTF containing each of the treatments overlayed with mineral oil. Embryos were imaged using a Zeiss LSM 800 confocal microscope (Carl Zeiss) with the incubation chamber set to 37 °C and 5% CO_2_ for live-cell imaging. A 20× objective, 405 nm laser (Ex 350 nm for GSH and 420 nm for Cys) and Zen Blue software (Carl Zeiss) were used for image capture. Embryos were analysed at the point where the nuclei were in focus. Microscope settings were reused between each condition and for each experiment.

### Analysis of confocal images

Images were analysed using Fiji by Image J to obtain the integrated density of fluorescence for GSH and Cys. The integrated fluorescence density of individual blastomeres was measured in three equidistant locations around the nuclei using a circle of equal size (Figure 2). The integrated intensities of the 3 areas for each blastomere were averaged and the corrected total cell fluorescence was then calculated using the following formula: Corrected total cell fluorescence = integrated density – (background fluorescence × mean area) [37]. Measurements for GSH and Cys were taken separately and reported individually.

**Figure 2.**
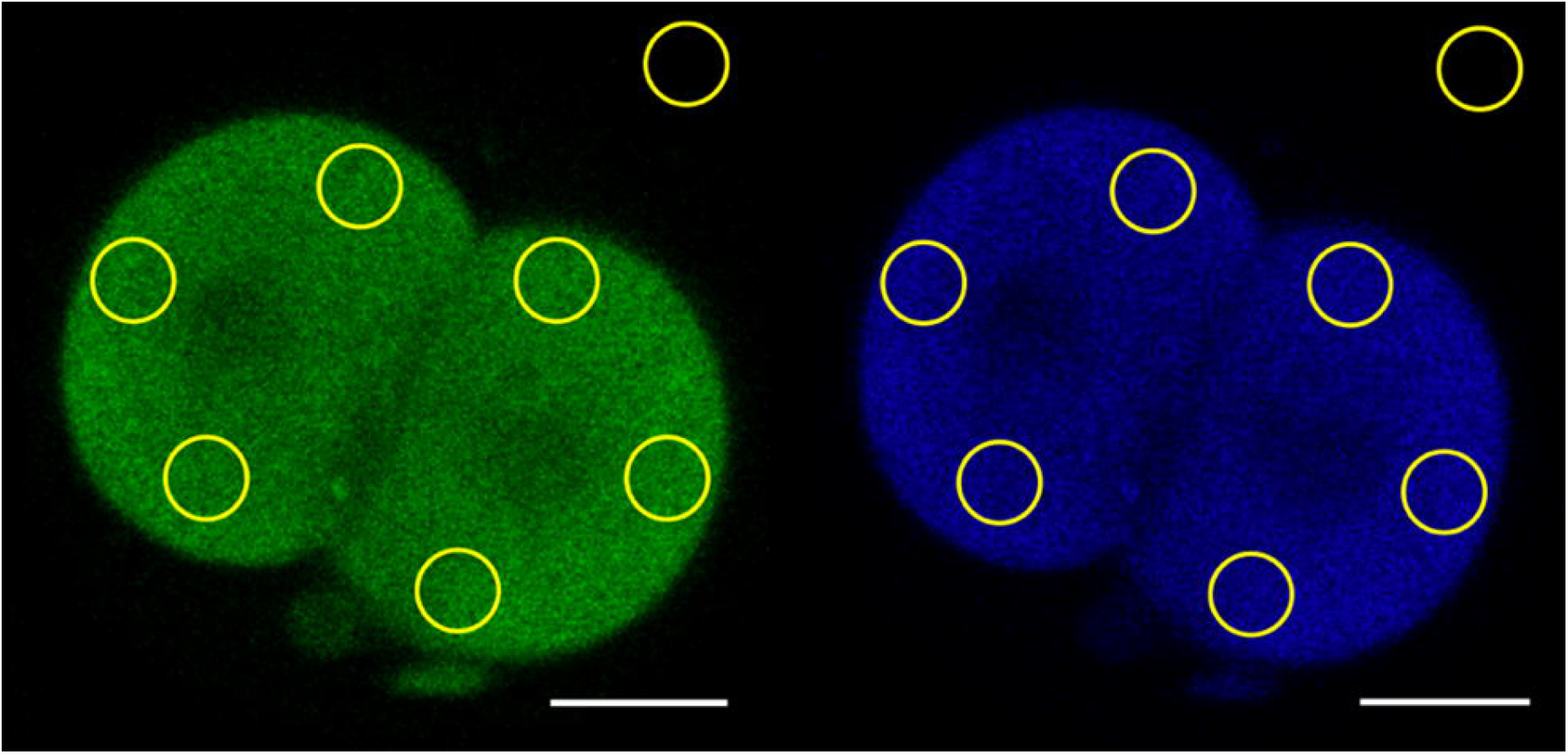
Data generation for live-cell (A) GSH and (B) cysteine imaging. Images taken on the Zeiss LSM 800 (Carl Zeiss) were opened using Fiji by Image J. The ellipse tool was used to measure three representative areas from each blastomere, equally spaced around the nucleus. Blastomeres were analysed individually to account for differences in fluorescence between individual cells. The average fluorescence of each embryo was then calculated. A background fluorescence measurement (ellipse in upper right corner) was taken to subtract from the blastomere fluorescence. Scale bar = 20 µm.

### Sample collection and extraction for LC-MS

Embryos were cultured in the presence or absence of 400 µM Pro to the zygote, 2-cell, 4-cell, 8-cell, morula or blastocyst stage as above and collected in groups of at least 20. Embryos were washed in PBS + PVA containing 2 mM NEM for 2 min to derivatise GSH content and transferred to microfuge tubes containing 100 µL HPLC-grade water prior to snap freezing in liquid nitrogen and storing at –80 ° C until extraction.

For measurement of GSH, samples were quenched and extracted using protocols adapted from Li et al. 2019 [38]. Briefly, frozen samples were thawed and extracted using 500 µL cold methanol with 0.15 µM s-hexylglutathione added as an internal standard. Samples were then centrifuged at 10,000 *g* for 15 min at 4 °C and the supernatant collected and stored at –80 °C for up to one week prior to concentration. Samples were dried using a vacuum concentrator (Eppendorf Vacfuge Plus) and then resuspended in 30 µL HPLC-grade water immediately before performing LC-MS. To allow quantification using a standard curve, standards of 20 mM stock of G-NEM (GSH with 2 mM NEM) were prepared and serially diluted to give concentrations ranging from 2 mM to 3.9 µM. A 10 µM stock of GSSG was prepared and serially diluted to give concentrations of 10 µM to 20 nM. The same extraction protocol was applied to standards as samples (see above) standards were resuspended in 60 µL HPLC water.

### Liquid chromatography-mass spectrometry (LC-MS)

All LC-MS was performed on a Shimadzu LC-20AD (Shimadzu, Kyoto, Japan) coupled to an AB SCIEX QTRAP 6500+ system (Applied Biosystems, Foster City, CA). Samples were held at 4 °C prior to injection. 5 µL sample was injected onto the column held at a temperature of 35 °C and a maximum pressure of 300 bar. Metabolites were separated using an InfinityLab Poroshell 120 Hilic Z column (Agilent). Elution was performed over 28 min using a multistep gradient of Buffer A (5% acetonitrile, 20 mM NH_4_OH and 20 mM NH_4_ acetate) and Buffer B (100% acetonitrile) with a constant flow rate of 200 µL/min. GSH and GSSG standards were run three times throughout the LC-MS run and water blanks were run as a negative control and to clean the column every 20 samples.

### Analysis of LC-MS data

MultiQuant (Sciex) was used to analyse peaks and baseline LC-MS data. Peaks for each metabolite in a sample were compared to the elution time of that metabolite in the standard. Area under the curves (AUCs) and retention time of individual peaks were obtained and transferred to Microsoft Excel. AUCs for metabolite standards were converted to mol/injection volume and the ratio of the peak area of metabolite standard to the peak area of the internal standard was calculated. Standard curves were plotted as the ratio of metabolite to internal standard ratio vs mol/injected volume and fitted by linear regression, from which the amount of metabolite in each sample was calculated. Mol/sample was then divided by the number of embryos in each sample and results reported as pmol/embryo.

### Statistics

GSH and Cys fluorescent images were analysed by Graphpad Prism using a oneway ANOVA with Tukey’s multiple comparisons test to compare fluorescence between stages. LC-MS data was analysed using Student’s *t*-test to compare the difference at each stage between embryos cultured in the presence or absence of Pro.

## Results

### Pro increases the amount of GSH in 2-cell, 4-cell and 8-cell embryos

To determine changes in intracellular concentrations of GSH, and its rate-limiting component Cys, the fluorescent probe 4F-2CN was used and embryos were imaged using confocal microscopy. Culturing 2-cell embryos in Pro for 24 h increased the fluorescence at 350 nm, and thus the GSH content (Figure 3A,B). However, culture with Pro did not affect the amount of Cys (Figure 3A,C). The culture of embryos with THFA alone, which inhibits POX and prevents the conversion of Pro to P5C, did not affect GSH amount and prevented the Pro-mediated increase in GSH. When embryos were cultured in the presence of NEM, which converts GSH to G-NEM, GSH fluorescence was reduced as expected. Similarly, culture in Cys increased Cys and GSH levels (Figure 3).

**Figure 3.**
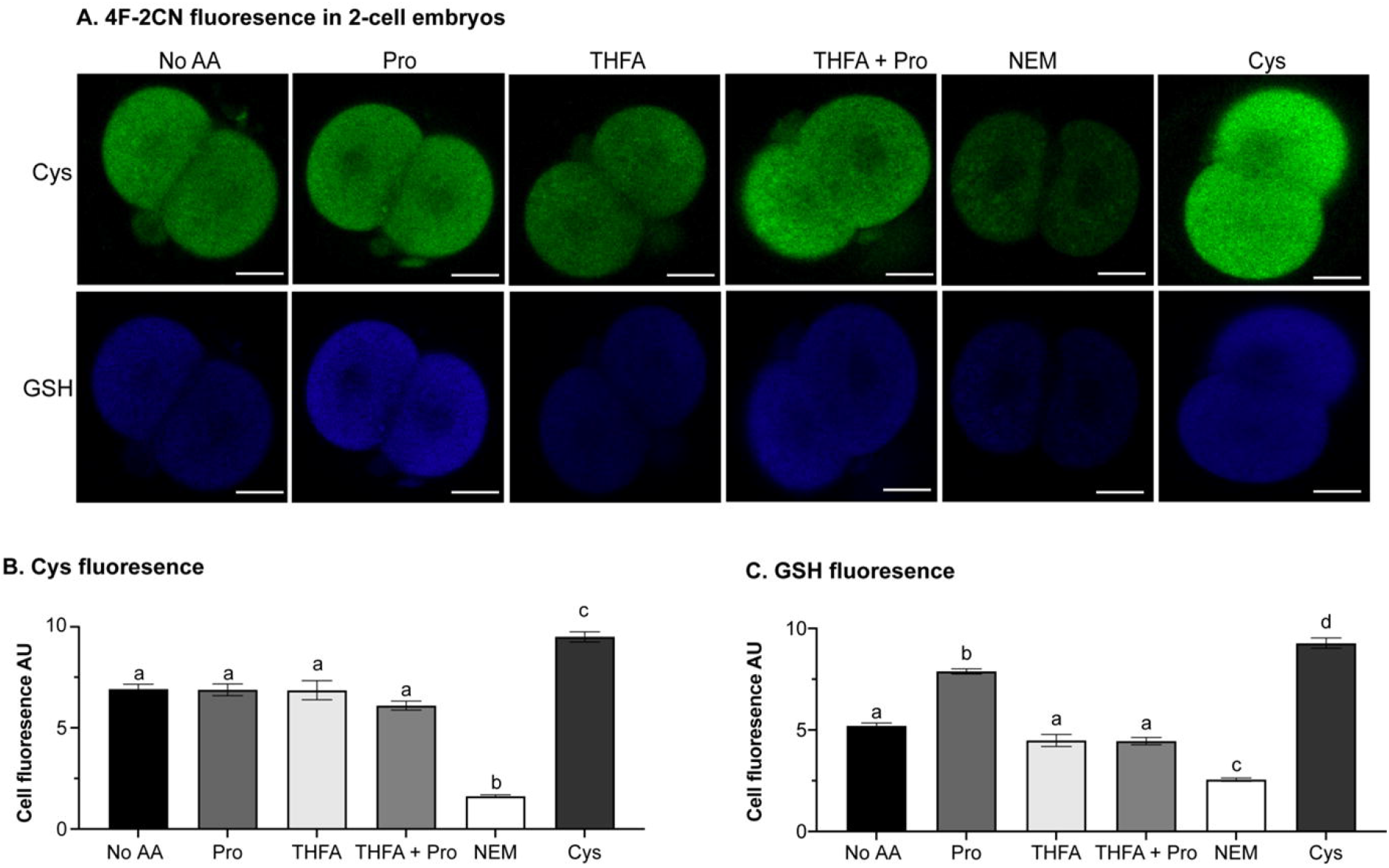
The effect of Pro on the amount of GSH and Cys in 2-cell embryos. Zygotes were cultured in ModHTF in the presence or absence of 400 µM Pro, THFA, or THFA + Pro for 24 h to the 2-cell stage. Embryos were loaded with 20 µM 4F-2CN and imaged using confocal microscopy. (A) Representative images of embryos in each condition, scale bar represents 20 µm. Cell fluorescence of (B) GSH (λ_ex_ 350 nm) and (C) Cys (λ_ex_ 420 nm). For positive and negative controls embryos were exposed to 2 mM NEM or 0.25 µM Cys, respectively, for 30 min prior to staining with 4F-2CN and imaging in media containing 2 mM NEM or 0.25 µM Cys. Error bars represent the mean ± SEM of 24-82 embryos obtained from at least 3 independent experiments. Data were analysed using a one-way ANOVA with a Tukey’s multiple comparisons test. Bars not sharing the same letter are significantly different (*P* <0.05).

Similar results were obtained for both 4- and 8-cell embryos: Pro increased GSH levels, THFA had no effect on its own but prevented the Pro-mediated increase (Figure 4A,B and 5A,B). NEM reduced GSH levels at both stages (Figure 4B and 5B).

**Figure 4.**
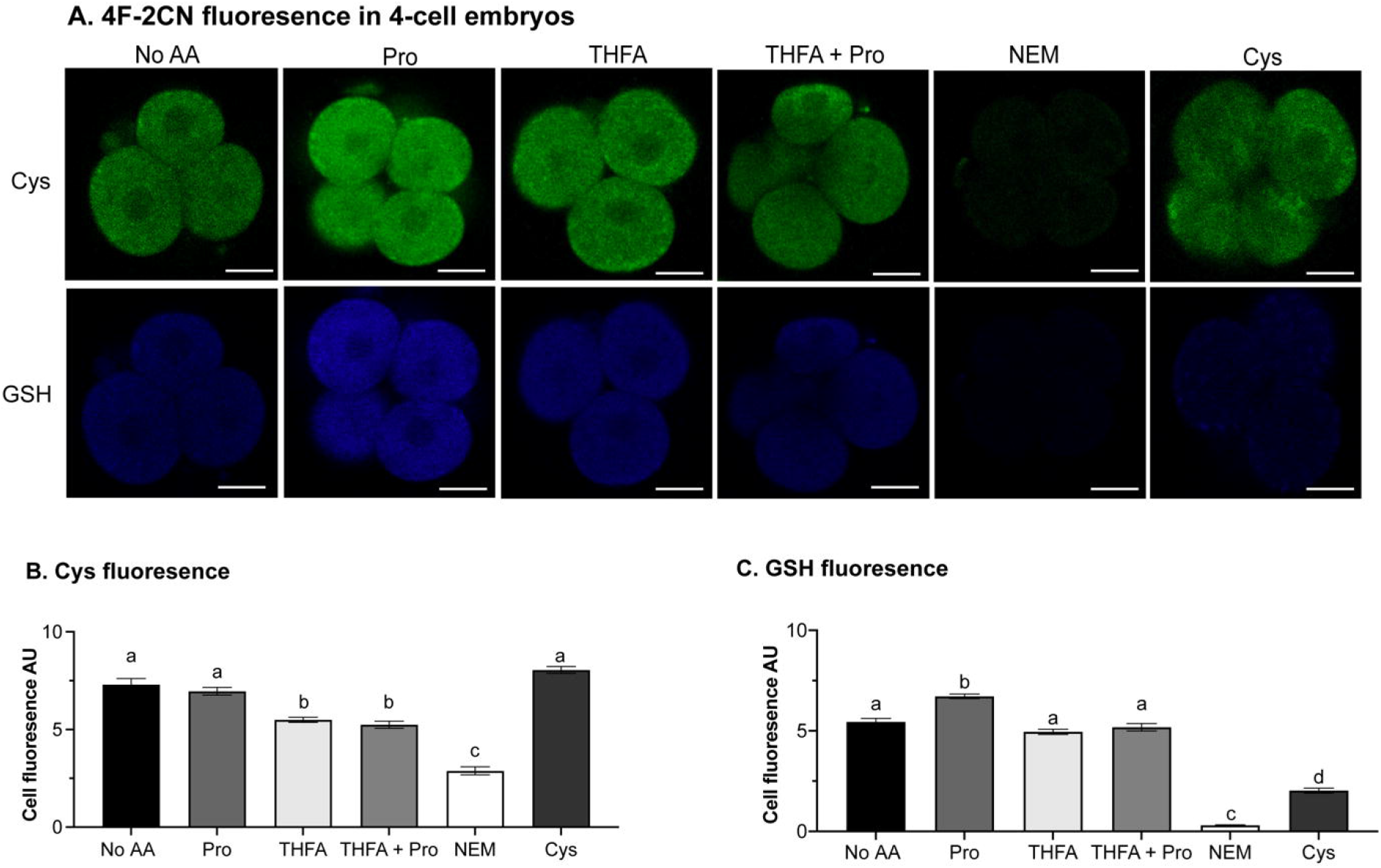
The effect of Pro on the GSH and cysteine content in 4-cell embryos. Zygotes were cultured in ModHTF in the presence or absence of 400 µM Pro, THFA or THFA + Pro for 36 h to the 4-cell stage. Embryos were loaded with 20 µM 4F-2CN and imaged using confocal microscopy. (A) Representative images of embryos in each condition, scale bar represents 20 µm. Cell fluorescence of (B) GSH (λ_ex_ 350 nm) and (C) Cys (λ_ex_ 420 nm). For positive and negative controls embryos were exposed to 2 mM NEM or 0.25 µM Cys, respectively, for 30 minutes prior to staining with 4F-2CN and imaging in media containing 2 mM NEM or 0.25 µM Cys. Error bars represent the mean ± SEM of 24-82 embryos obtained from at least 3 independent experiments. Data were analysed using a one-way ANOVA with a Tukey’s multiple comparisons test. Bars not sharing the same letter are significantly different (*P* <0.05).

**Figure 5.**
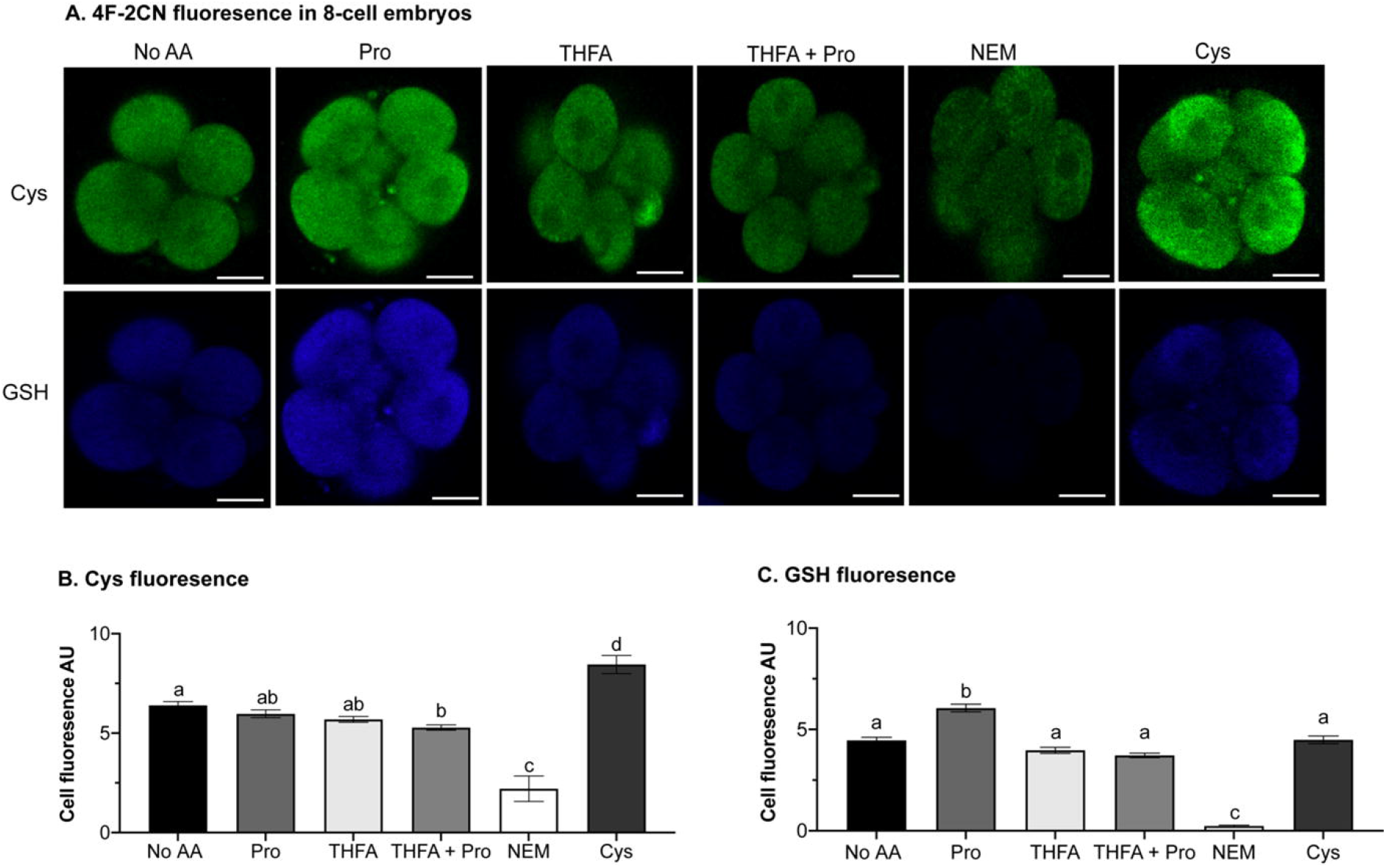
The effect of Pro on the GSH and cysteine content in 8-cell embryos. Zygotes were cultured in ModHTF in the presence or absence of 400 µM Pro, THFA or THFA + Pro for 48h to the 8-cell stage. Embryos were loaded with 20 µM 4F-2CN and imaged using confocal microscopy. (A) Representative images of embryos in each condition, scale bar represents 20 µm. Cell fluorescence of (B) GSH (λ_ex_ 350 nm) and (C) Cys (λ_ex_ 420 nm). For positive and negative controls embryos were exposed to 2 mM NEM or 0.25 µM cysteine, respectively, for 30 minutes prior to staining with 4F-2CN and imaging in media containing 2 mM N-ethylaemide (NEM) or 0.25 µM Cys. Error bars represent the mean ± SEM of 24-82 blastomeres obtained from at least 3 independent experiments. Data were analysed using a one-way ANOVA with a Tukey’s multiple comparisons test. Bars not sharing the same letter are significantly different (*P* <0.05).

Addition of Cys decreased, rather than increased, GSH levels in 4-cell embryos. However, Cys had no effect on GSH levels in 8-cell embryos. Addition of Cys had no effect on the amount of Cys in 4-cell embryos but did increase Cys levels in 8-cell embryos (Figure 4C and 5C).

### Embryo culture in Pro increases the amount of GSH throughout preimplantation development

As the results of the previous experiment using live-cell imaging indicated that Pro increases GSH levels, we used LC-MS to quantify this increase at each preimplantation stage from zygote to blastocyst as well as the ratio of GSH to GSSG.

Culture in Pro increased the amount of GSH in the embryo at all stages of pre-implantation development (Figure 6A) but resulted in little or no change in GSSG amounts (Figure 6B). However, the GSH:GSSG ratio was increased in the presence of Pro at all stages, indicating reduced oxidative stress (Figure 6C) [40]

**Figure 6.**
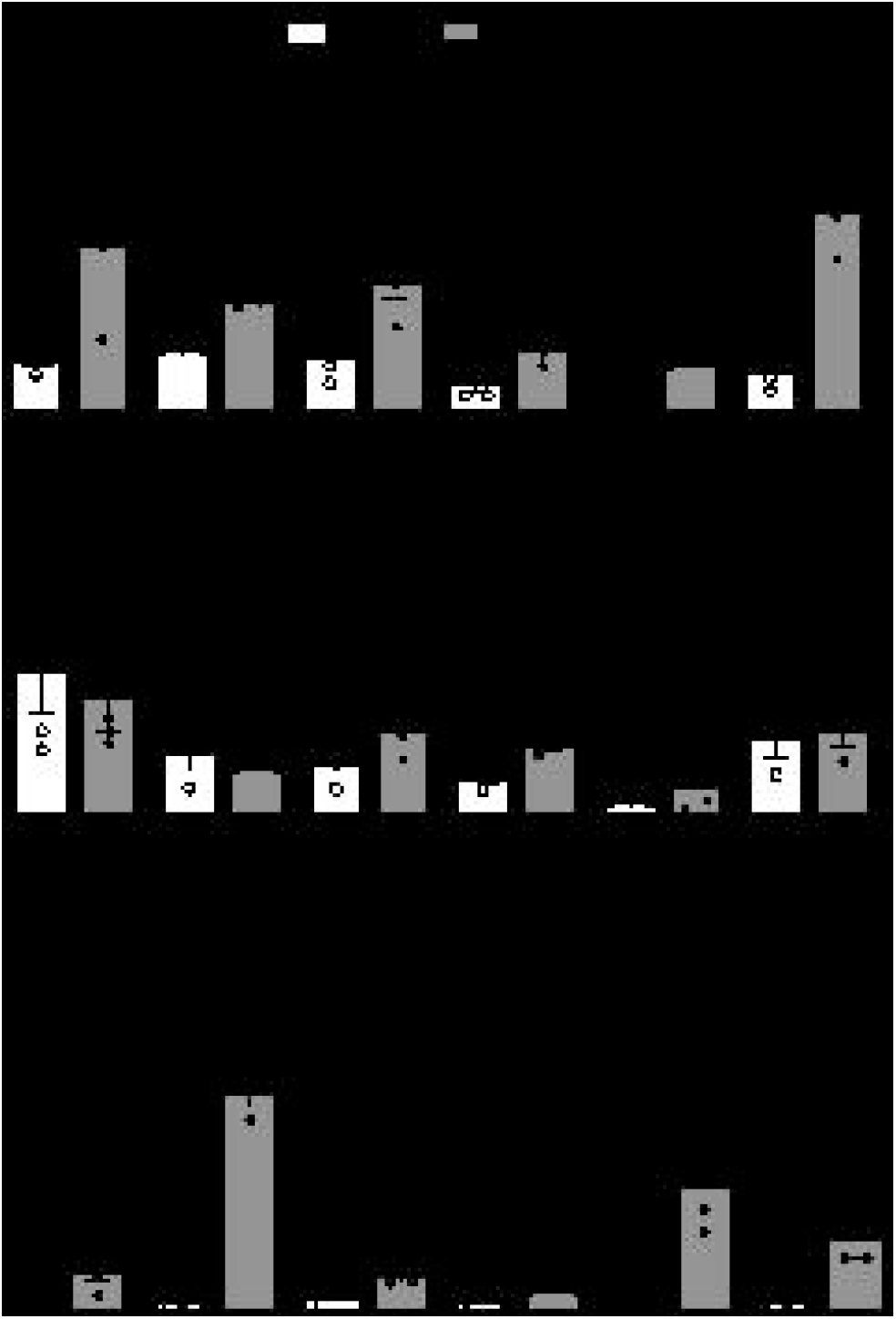
Intracellular amounts of (A) GSH and (B) GSSG amounts, and (C) GSH:GSSG detected by LC-MS in mouse preimplantation embryos cultured in ModHTF in the absence (No AA) or presence of 400 µM Pro to the zygote (3 h), 2-cell (24 h), 4-cell (34 h), 8-cell (48 h), morula (66 h) and blastocyst (78 h) stages. To quantify GSH, embryos were treated with 2 mM NEM for 2 min to convert GSH to G-NEM. Error bars represents the mean ± SEM of 3-4 samples. Each point represents a sample of 20-30 embryos. Data were analysed using a t-test * *P* <0.05, ** *P* <0.001, \*\*\**P* <0.0001.

## Discussion

This study demonstrates that the addition of Pro to culture medium from the zygote stage increases the GSH content in 2-cell, 4-cell and 8-cell stage embryos, as indicated by live-cell fluorescence imaging with 4F-2CN, and at all stages of preimplantation development as shown with LC-MS. Furthermore, Pro increases the GSH:GSSG ratio at all stages.

GSH levels increase *in vivo* as the oocyte matures from germinal vesicle to metaphase II stages [41]. MII oocytes have the highest GSH content (7 pmol/oocyte) which appears to protect it against the oxidative burst associated with sperm capacitation and fertilisation [42], after which GSH content decreases [16]. Results presented here are concordant with these results with a reduction in GSH between the zygote and morula. This is further supported by the qPCR results where *Gpx1* and *Gpx4* expression is decreased in the morula and the blastocyst stages of development suggesting less GSH production. Similarly, Gss, which catalyses the addition of Gly to γ-glutamylcysteine for GSH synthesis, has reduced expression after the 8-cell stage. Our results found that in the blastocyst there was less GSH than the morula (*P* =0.0014). These discrepancies in the blastocyst results cannot be closely analysed as the findings reviewed by Hansen and Harris (2015) were not published, thus, differences in methodology, including the stage of blastocyst formation or choice of culture media, cannot be compared [42]. However, it is possible that blastocysts have lower GSH content, for a number of reasons. Firstly, *in vivo* as the blastocyst moves into the uterus the oxygen concentration decreases from 5-7% in the oviduct to 2% in the uterus, thus decreasing oxidative stress and the requirement for GSH [43]. Although our *in vitro* experiments did not have dynamic oxygen concentrations and were completed in atmospheric oxygen levels, it is possible that blastocysts have naturally decrease GSH due to the oxygen levels they are exposed to *in vivo*. Further, embryos undergo a burst of ROS production during blastocyst hatching which is required for correct initiation of hatching from the zona pellucida. The potent antioxidant superoxide dismutase decreases in concentration by up to 50% between pre-hatching and peri-hatching blastocyst stage, to allow this oxidative burst to occur, therefore it is possible a similar mechanism occurs with GSH with the concentration decreasing to allow for blastocyst hatching. [44].

GSH is critical for embryo development to keep oxidative stress balanced. When GSH stores are depleted in embryos cultured *in vitro* using NEM (the same compound we use to quench GSH for measurement in LC-MS),,embryos are unable to produce GSH and reach the morula and blastocyst stages thus indicating the criticality of GSH for normal embryo development to ensue [16]. This occurs even when Cys, the rate limiting component of GSH synthesis, is added to culture media to facilitate GSH recovery. As Pro is metabolised to Glu a precursor of GSH, it became a feasible avenue to investigate the mechanism by which Pro was acting to improve embryo development.

LC-MS data indicated that Pro increased GSH levels across all stages of development which supports the theory that one way Pro improves embryo development by is increasing GSH stores. These results support similar findings in mouse oocytes, which when cultured in 500 µM Pro increased GSH content and decreased ROS [30], and in porcine trophectoderm cells cultured with 500 µM Pro showing increased total intracellular GSH levels [45]. Our results also show that all stages of preimplantation embryo development are capable of synthesising GSH.

The Pro-mediated improvement in embryo development begins at the at the late 2-cell stage [27], thus the changes in GSH and Cys levels at the 2-cell to 8-cell stages were studied to determine if there was a relationship between GSH levels and the effect of Pro. However, mass spectrometry data identified that GSH levels and the GSH:GSSG were increased by Pro at all stages throughout preimplantation embryo development.

In order to elucidate how Pro increases GSH, the inhibitor THFA which blocks POX, was used in these experiments. THFA inhibits the conversion of Pro to P5C, preventing continuation of the downstream Pro metabolic pathway [50]. Culture of embryos with Pro + THFA at the 2-cell, 4-cell and 8-cell stages of development universally prevented Pro-mediated increases in GSH, therefore Pro metabolism by POX is required for Pro to increase GSH. Culture with THFA alone did not change GSH amounts. Unexpectedly, THFA did not decrease GSH levels, this suggests that endogenous Pro typically does not contribute greatly to GSH production, this may because *in vivo* Pro is at much lower concentrations in oviductal (0.14 mM [52] and uterine fluids, as well as intracellularly in stem cells, than other amino acids including other GSH precursors such as glutamate and glutamine 1 mM in oviductal fluid [53] and potentially is also lower in preimplantation embryos, thus it is only when exogenous Pro is added to culture media that Pro is used that Pro is used as a substrate for GSH production. This is similarly seen in culture of retinal pigment epithelial (RPE) cells with THFA to inhibit Pro oxidation decreased GSH concentration, indicating the important role that Pro metabolism plays in the increase of GSH [54].

The GSH:GSSG ratio is a parameter of oxidative stress which is more representative of a cells true oxidative state than GSH measures alone. As cells are exposed to oxidative stress, GSSG accumulates, with more of the total glutathione pool being stored in this oxidised form (GSSG) [16,40]. In a resting cell, the expected ratio of GSH:GSSG is 100:1. Exposure to oxidative stress in the form of ROS shifts this balance; in an oxidative stress state, the ratio can shift to 10:1 or even 1:1 [40]. This agrees with the current LC-MS results, in embryos cultured in ModHTF lacking antioxidants, the ratio of GSH:GSSG ranged from 7:1 to 19:1 (Figure 4.6). By contrast, embryos cultured in Pro had GSH:GSSG ratios ranging from 51:1 to 312:1. These data provide evidence that oxidative stress is substantially less in preimplantation embryo development when cultured in the presence of Pro as demonstrated by the increased amount of GSH in the cell in the embryo compared to GSSG. In the glutathione cycle, oxidative stress is mitigated by increasing GSH levels with efficient recycling of GSSG by glutathione reductase to GSH. This means that GSSG to combat oxidative stress GSSG levels typically remain constant while GSH levels are increased to combat oxidative stress [55,56].

Despite the increases observed in GSH, live-cell imaging with 4F-2CN showed that culture with Pro did not change Cys. This is unsurprising, as Pro is not metabolized into constituents that contribute to Cys production. Cys is primarily formed from the metabolism of Ser and Met [57]. Although Pro increases GSH, it was expected that Cys would decrease, as it is required for GSH synthesis. However, no significant decrease in Cys levels was observed in embryos cultured with Pro. This may be due to Cys synthesis from other pathways, including Ser metabolism and GSH degradation.

In post-compaction embryos, Ser has one of the highest amino acid consumption rates (~1 pmol/embryo/hour), while Cys has a one of the highest production rates in spent culture media (~0.5 pmol/embryo/hour). Although Ser is not included in the culture media, intracellular Ser stores may support Cys synthesis. A decrease in intracellular Ser levels is observed between the 2-cell and 4 to 8-cell stages, indicating increased Ser utilization [58]. These findings suggest Cys levels can be quickly replenished, preventing a drop in Cys due to Pro metabolism.

Culture in Cys reduced GSH content in the 4-cell embryo and did not affect GSH content in the 8-cell embryo. As Cys is a precursor to GSH, an increase in GSH, like seen here in 2-cell embryos, was expected. As Cys did not increase Cys fluorescence at the 4-cell stage, it is possible that there was a greater rate of consumption of Cys than rate of production. The addition β-mercaptoethanol with Cys may promote uptake of Cys from the media, preventing the autooxidation between cysteine and cystine [59]. This does not however explain why Cys reduces GSH levels to lower than *in vitro* developed embryos without AAs. One potential reason is that culture in Cys may change gene expression or enzyme activity preventing GSH synthesis or conversion to GSSG. Culture of cardiomyocytes with Cys increased Gpx1 activity [60]. Gpx1 converts GSH to GSSG, which is not captured by looking at GSH alone which is done with 4F-2CN in the present study, thus potentially a similar phenomenon is occurring at the 4-cell and 8-cell stages. PCR and enzyme activity and LC-MS to look at the changes in the GSH:GSSG ratio would confirm this.

The expression and activity of numerous AA transporters change throughout the different embryonic stages resulting in stage-dependent uptake of different AAs. Pro is known to be taken up in all stages of development, being taken up by transporters B^0^AT1, GLYT1, PAT1/2 throughout development [23,61–64]. Exposure of 2- and 8-cell but not 4-cell embryos to Cys increased intracellular Cys suggesting uptake by AA transporters at the 2- and 8-cell stages. The AA transporters for Cys in preimplantation embryos have not been characterised, although Cys can be taken up by a number of transporters, including ASCT2, LAT2, xCT and b^0,+^AT. However, the expression or activity of these transporters at the 4-cell stage is not known [22,65]. There is a transient rise in xCT activity between oocytes and zygotes, however, no other changes in transporter activity have been identified [66]. Uptake experiments using radiolabelled Cys would allow Cys uptake throughout development to be determined.

*In vitro* culture increases ROS, which has detrimental effects on preimplantation embryos. Combating excessive ROS production has been a key focus of assisted reproduction and embryo development research [29,67–71]. As discussed, GSH is a potent antioxidant, and although in bovine embryos addition of 3 mM GSH did improve blastocyst development addition of GSH to culture media itself has inconsistent results and is found to be less effective than addition of its constituents [16,72,73]. This may be due to the fact, GSH cannot be taken up into the embryo as a tripeptide and hence is split into individual amino acids prior to uptake, as discussed, the uptake of AAs changes with developmental regulation of AA transporters [74,75]. Therefore, adding GSH itself to culture media is not a feasible solution. This reinforces the idea that adding Pro, which can be taken up throughout embryo development may be a way to reduce ROS. This study, as well as the aforementioned studies on boar sperm, porcine trophectoderm cells and RPE cells, all support the addition of Pro to stimulate GSH production. GSH production is also stimulated with the addition of cysteine, cysteamine and β-mercaptoethanol [59].

This study, as well as previous studies have identified multiple mechanisms by which Pro reduces ROS, including but not limited to, direct ROS scavenging whereby an electron is donated from the secondary amine group of Pro to quench free radicals [76], although previous studies suggest this is not occurring in preimplantation embryos as inhibition of POX prevented Pro reducing ROS [29]. Pro also reduces mitochondrial activity via direct interaction with the electron transport chain [29,77] and activates the mTOR pathway and Erk and Akt pathway [27,78]. Finally, the current study shows that Pro increases GSH synthesis and the GSH:GSSG ratio protecting the embryo against stage specific variations in oxidative stress and to ensure the GSH: GSSG ratio remains in a tight physiological range. Although the exact reason there are so many mechanisms by which Pro appears to be acting to improve embryo development is not known, it is likely that a dynamic complex system working simultaneously to achieve the final outcome of improved embryo development.

